# The individual vulnerability to develop compulsive adjunctive behavior is associated with the recruitment of activity-regulated cytoskeleton-associated protein (Arc) within the Locus Coeruleus

**DOI:** 10.1101/2022.09.13.507749

**Authors:** Clara Velazquez-Sanchez, Leila Muresan, David Belin

**Author notes:** **Corresponding author:** Professor David Belin, Department of Psychology, University of Cambridge, Downing St., Cambridge CB2 3EB, UK.

## Abstract

Some compulsive disorders have been considered to stem from the loss of control over coping strategies, such as displacement. However, the cellular mechanisms involved in the acquisition of coping behaviors and their ensuing compulsive manifestation in vulnerable individuals have not been elucidated. Considering the role of the locus coeruleus (LC) noradrenaline dependent system in stress and related excessive behaviors, we hypothesised that neuroplastic changes in the LC may be involved in the acquisition of an adjunctive polydipsic water drinking, a prototypical displacement behavior, and the subsequent development of compulsion in vulnerable individuals. Thus, male Sprague Dawley rats were characterised for their tendency, or not, to develop compulsive polydipsic drinking in a schedule-induced polydipsia (SIP) procedure before their fresh brains were harvested. A new quantification tool for RNAscope assays revealed that the development of compulsive adjunctive behavior was associated with a low mRNA copy number of the plasticity marker Arc in the LC which appeared to be driven by specific adaptations in an ensemble of tyrosine hydroxylase (TH)+, zif268-neurons. This ensemble was specifically engaged by the expression of compulsive adjunctive behavior, not by stress, because its functional recruitment was not observed in individuals that no longer had access to the water bottle before sacrifice while it consistently correlated with the levels of polydipsic water drinking only when it had become compulsive. Together these findings suggest that downregulation of Arc mRNA levels in a population of a TH+zif268-LC neurons represents a signature of the tendency to develop compulsive coping behaviors.

## Introduction

Failure adaptively to regulate emotions, such as coping with stress, has been associated with an individual vulnerability to develop several neuropsychiatric disorders, such as anxiety and depression, PTSD as well as Impulsive/Compulsive Spectrum Disorders including Obsessive Compulsive and substance use disorder [1–5]. Thus, the emergence of compulsion, which characterises the excessive and persistent nature of several compulsive disorders [6] has been suggested to stem from a loss of control over coping strategies [7–16], such as displacement behaviors, which initially aim to decrease negative affect or stress in adverse situations but progressively become habitual and rigid [17,18].

Across species, adjunctive behaviors [19] represent a form of displacement activity [20,21] in which individuals engage to decrease stress [20,22–24]. One such adjunctive anxiolytic response, schedule-induced polydipsia (SIP) [20,21,25–29], a non-regulatory polydipsic water drinking in the face of intermittent food delivery in food-restricted animals [29–34] decreases the levels of stress-related hormones [21,25–27,35–39] and autonomic nervous system responses to stress [39], a decrement that is not observed when individuals do not have access to water [25].

Like in humans, the majority of individuals who engage in coping displacement tend, at the population level, to maintain relative control over their adjunctive response [40–44]. However, some vulnerable individuals, characterized, for instance by a high impulsivity trait [43], lose control over their polydipsic intake which becomes excessive and inflexible [40–46]. The psychological and neural basis of this individual vulnerability to develop compulsive adjunctive behaviors has not been fully elucidated. While research into the neural systems basis of adjunctive behaviors, including their compulsive manifestation, has primarily focused on dopaminergic and serotoninergic mechanisms within the limbic corticostriatal circuitry [45–47], increasing evidence supports a role of the noradrenergic system [44,48] otherwise long known for its contribution to stress and anxiety [49]. Thus, not only has the acquisition of different coping responses been associated with specific changes in the neural mechanisms that control the locus coeruleus (LC) noradrenaline (NA) stress response system [50] and associated coping [51–54], but the integrity or functionality of the LC has also been suggested to be necessary for the expression of polydipsic adjunctive water intake at the population level [55].

In addition, the noradrenaline reuptake inhibitor, atomoxetine, prevented the development of compulsive polydipsic behavior in vulnerable rats characterised for their high impulsivity trait [44]. Together with the evidence that central noradrenergic mechanisms mediate the emergence of stress-induced repetitive behaviors [56], these observations suggest that specific adaptations taking place in the LC may contribute to the emergence of compulsive adjunctive behaviors in vulnerable individuals. However, the LC is a highly functionally and cytoarchitecturally heterogeneous structure [57,58] with subdivisions or particular neuronal ensembles [59,60] or microcircuits [61] involved in different aspects of behavior [62–64], including stress-related mechanisms [65].

At the cellular level, the response of the LC-NA system to a variety of stressful situations including restrain, exposure to mild electric shocks or social stress has been associated with the recruitment of immediate early genes (IEGs) [66,67] including activity or plasticity-related transcription factors, such as c-fos or zif268, respectively, and effectors, such as activity-related cytoskeleton-associated protein (Arc), which instead directly influence cellular processes other than gene transcription [68]. Thus, exposure to novelty, anxiogenic drugs, repeated episodes of restraint stress or social stress result in an increase in c-fos in the LC [50,69–74]. However, c-fos mRNA levels tends to decrease over repeated exposure to stressful situations [75,76] which precludes its use as an ensemble marker in the context of the present study. This is not the case of Arc [77], an effector of BDNF, glutamatergic, dopaminergic and serotoninergic signalling, involved in learning-associated synaptic and dendritic plasticity [78,79] which mRNA levels in the LC have been shown, using standard *in-situ* hybridisation not to respond to acute stress, but instead to situations of adaptation to chronic challenges [80].

Thus, here we investigated whether the emergence of compulsive adjunctive behavior was associated with the recruitment of a neuronal ensemble in the LC that is characterised by a selective engagement or Arc-dependent cellular plasticity. Because Arc transcription is under the control of zif268, even though their mRNA levels do not necessarily correlate [81], we sought to determine whether any potential Arc-dependent ensemble was specifically engaging Arc or whether it was also reflecting the activation of zif268. For this we developed a new RNAscope multiplex assay quantification method in order to investigate the mRNA copies in genetically identified cellular ensembles in the LC of rats with controlled or compulsive adjunctive behavior sacrificed forty-five minutes after a challenge SIP session during which they expressed their polydipsic drinking behavior or were prevented from doing so.

## Methods and Materials

### Subjects

Forty-eight male Sprague Dawley rats (Charles River, UK) weighing approximately 300 g at the start of the experiment were single-housed under a reversed 12h light/dark chain (lights off at 7:00 AM). After a week of habituation to the vivarium, rats were food restricted to gradually reach 80% of their theoretical free-feeding body weight before starting behavioral training. Water was always available *ad libitum.* Experiments were performed 6-7 days/week between 8AM-5PM. All experimental protocols were conducted under the project license 70/8072 held by David Belin in accordance with the regulatory requirement of the UK Animals (Scientific Procedures) Act 1986, amendment regulations 2012, following ethical review by the University of Cambridge Animal Welfare and Ethical Review Body (AWERB).

### Timeline of the experiments

As illustrated in **Figure 1A**, after one week of habituation to the animal facility, rats were progressively food restricted to 80% of their theoretical free-feeding body weight. They were accustomed to the SIP context over two habituation sessions during which their regulatory water intake was measured and then trained in a SIP procedure for 21 daily sessions. On day 22, rats underwent a 60 min challenge session with or without the opportunity to express their adjunctive behavior, forty-five minutes after which they were sacrificed, and their fresh brains harvested and properly stored for subsequent RNAscope assays.

**Figure 1:**
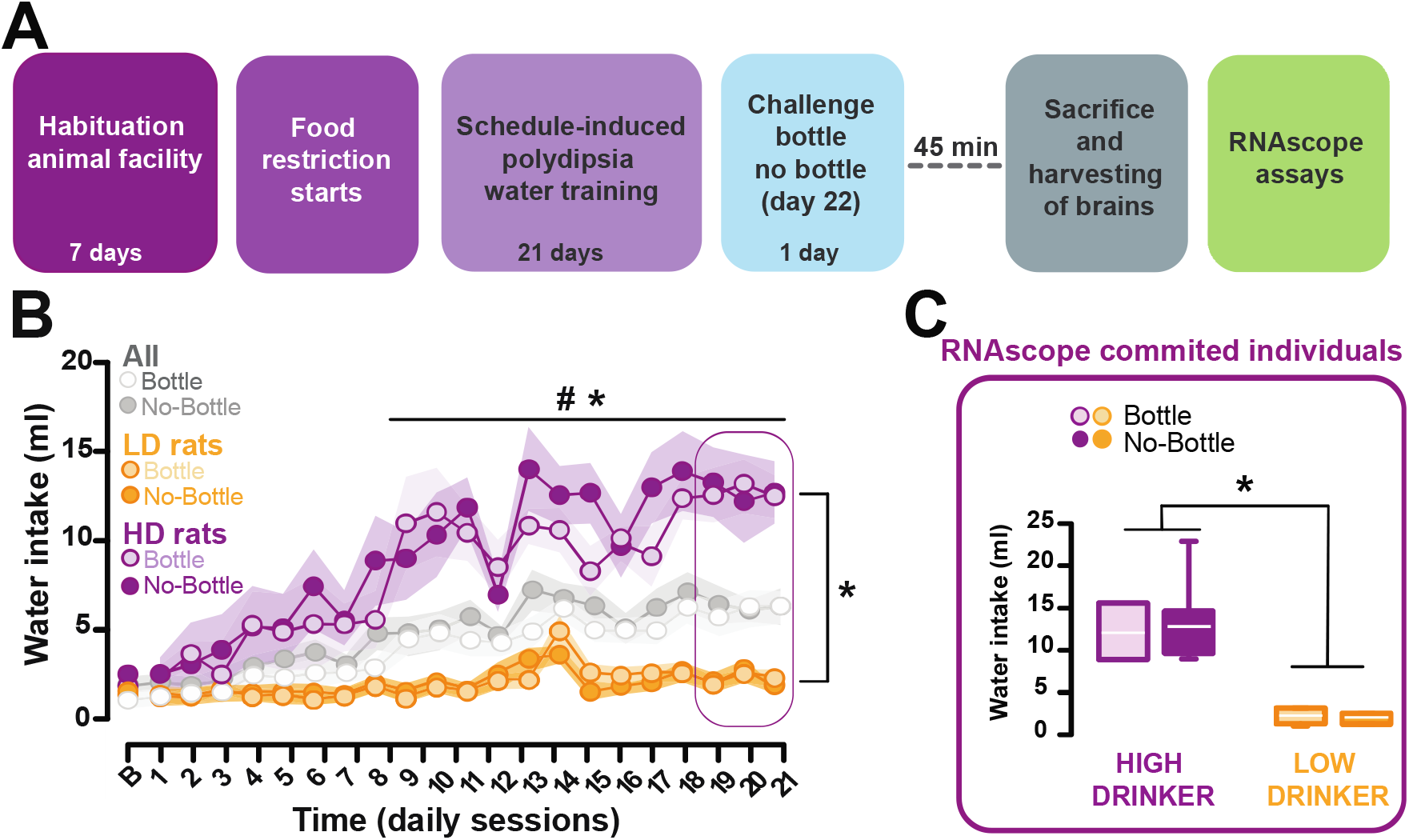
General scientific strategy and timeline of the experiments involved in the characterisation of the individual vulnerability to develop compulsive adjunctive behavior. **A)** After a week of habituation to the vivarium forty-eight male Sprague Dawley rats were food restricted to gradually reach 80% of their theoretical free-feeding body weight. Then, rats were trained under a FT60s Scheduled-Induced polydipsia (SIP) procedure for twenty-one 1h daily sessions. High drinkers (HD) and Low drinkers (LD) rats were selected in the upper and lower quartile of the population, respectively, based on their average water intake during the last 3 sessions. Forty-five minutes after a last 1h session under FT60 during which half the population was prevented from expressing their adjunctive response by removal of the water bottle rats were sacrificed, their brains collected and further processed for RNAscope. **B)** While at the population level individuals acquired an adjunctive polydipsic water drinking behavior under a FT60s schedule of food delivery [main effect of session: session *F*_20,320_ = 14.03, *p* ≤ 0.001, pη^2^= 0.46], some vulnerable HD individuals progressively developed an excessive polydipsic water intake [main effect phenotype: *F*_1,16_ = 32.76, *p* ≤ 0.001, pη^2^= 0.67, and phenotype x session interaction *F*_20,320_ = 8.01, *p* ≤ 0.001, pη^2^= 0.33]. HD rats started excessively to consume water from session 8 onwards, a point at which they showed a level of drinking that was both higher than that related to homeostatic needs as measured at baseline (all *p*s ≤ 0.001) and different to the water intake displayed by the LD rats (all *p*s ≤ 0.05). **C)** HD and LD rats that were allocated to the bottle or non-bottle condition at test displayed different levels of adjunctive drinking behavior measured over the last 3 SIP sessions [main effect phenotype: *F*_1,16_ = 45.76, *p* ≤ 0.001, pη^2^= 0.74]. However, no differences were observed between the bottle or non-bottle conditions at test in HD or LD rats [phenotype x session interaction *F*_1,16_ = 0.058, *p*= 0.81, pη^2^= 0.0036]. Ten representative individuals of the HD and LD groups were selected to carry out the RNAscope assays. * Different from LD rats; *p*s ≤ 0.05. # HD rats different from baseline; *p*s ≤ 0.001).

### Apparatus

The SIP procedure was carried out as previously described [44,82] in 12 operant chambers located in ventilated and sound-attenuating cubicles (Med Associates, St. Albans, VT) controlled by MedPC software (Med Associates Inc., Ltd). Each operant chamber was made of aluminium and transparent acrylic plastic with a stainless-steel grid floor (24 x 25.4 x 26.7 cm) and was equipped with a house light (3-W), a food tray (magazine), installed at the centre of the front wall, and a bottle from which a stainless-steel sipper tube delivered water into a receptacle placed in a magazine on the wall opposite the food magazine.

### Schedule-induced polydipsia (SIP)

The day before the first habituation session rats were exposed in their home cages to the same 45 mg food pellets used in the SIP procedure in order to avoid any neophobia. On a first habituation session, rats were exposed for one hour to the operant chambers and were given access to water and 60 food pellets (45mg, *TestDiet, USA)* that were previously placed in the food magazine. On the second 1-hour habituation session, rats were exposed to a random interval (RT-60 seconds) schedule of pellet food delivery in order to ensure their regulatory water intake was measured over a 60 min period while they learnt that food was being dispensed in the magazine.

The SIP procedure was based on fixed-time 60-second schedule of food delivery, conditions previously shown to result in marked individual differences in the propensity to develop compulsive adjunctive drinking behavior [40,44,82]. Twenty-four hours after the baseline session rats underwent 21 of these FT-60s SIP sessions [44,83]. Three hundred ml bottles were filled daily with fresh tap water, weighted, and placed into the operant boxes immediately before the start of each session. House lights were switched on at the beginning and switched off at the end of each session. The total volume of water consumed during the session was calculated daily as the difference between the bottle weight before and after the session.

Water intake over the last 3 sessions was used to identify rats in the upper (High drinkers, HD) and lower quartile of the population (Low drinker, LD), as previously described [44,84].

Subsequently, in order to establish whether cellular correlates of the tendency to develop compulsive polydipsic behavior were attributable specifically to the expression of the anxiolytic adjunctive response or instead to the distress induced by the procedure, on day 22 rats underwent one 1h challenge SIP session during which they had access to the bottle of water, and could express their adjunctive response or not, thereby being prevented from engaging in their well-established coping habit [41], which we speculated should result in negative urgency [85] associated with heightened stress and frustration [23,86].

### Histology

45-min after the challenge session, animals were briefly anesthetized with isoflurane (<30s), decapitated and their fresh brains harvested, snap frozen at −40°C in isopentane (Sigma-Aldrich) and stored at −80°C, as previously described [85]. Brains were then processed using a cryostat (Leica Microsystems) into 12 μm thick coronal sections collected on Superfrost gelatine-coated slides (Fisher Scientific) and stored at −80°C until they were processed for multiplex RNAscope® in situ hybridization.

### RNAscope® in situ hybridization assay

RNAscope was performed according to the manufacturer’s instructions for fresh frozen tissue using the RNAscope Multiplex Fluorescent Reagent Kit (Advanced Cell Diagnostics). Brain sections were first fixed in chilled 10% NBF for 30 min on ice, rinsed 3 times in PBS and dehydrated in increasing concentrations of ethanol (50, 70, 100 and 100%). Slides were then kept in fresh 100% ethanol at −20°C overnight. The following day, sections were air dried and heated at 37°C on a hot plate for 20 min to prevent their detachment from the slides. Next, a hydrophobic barrier was drawn around each section to avoid the loss of the reagents during the assay. Sections were treated with Protease IV for 20 min at room temperature and then rinsed 3 x 5 min with PBS prior to being incubated with the target probes in a HybEZ oven for 2h at 40°C.

Each probe consists of a unique oligonucleotide mixture designed to bind to a specific target RNA, which, for this study, was Early Growth Response 1 (Egr1/zif268 probe) [GenBank accession number NM_012551.2, target nt region 162-1333], Tyrosine Hydroxylase (TH probe) [GenBank accession number NM_012740.3, target nt region 422-1403], Glial Fibrillary Acidic Protein (GFAP probe) [GenBank accession number NM_017009.2, target nt region 1539-2534], Activity-Regulated Cytoskeleton-Associated Protein (Arc probe) [GenBank accession number NM_019361.1, target nt region 269-1148]. Due to a limitation to 3 fluorescence microscopy filters, two different complementary assays were performed combining the target probes with different colour channels as follows: Egr1/zif268 (channel1)-TH (channel 2)-Arc (Channel4), and Egr1/zif268 (channel1)-GFAP (channel 2)-Arc (Channel4).

Following the 2h-incubation with the target probes, sections were incubated with the preamplifier and amplifier probes (AMP1, 40°C for 30 min; AMP2, 40°C for 15 min; AMP3, 40°C for 30 min) and washed with washing buffer in between each incubation step for 3x 5min. Then sections were incubated with the fluorescence labelled probe (AMP4 AltB-FL) to detect the three triple combination channels in orange (Alexa Fluor 550 nm), green (Alexa Fluor 488 nm) and far red (Alexa Fluor 647 nm), respectively. The sections were rinsed in washing buffer and incubated with DAPI for 20s before being coverslipped with Fluoroshield mounting medium (Abcam, ab104135).

### RNAscope® in situ hybridization imaging and quantification

Images for quantitative RNAscope analysis were captured with a Zeiss Axio Imager M2 equipped with an AxioCam MRm camera (Oberkochen, Germany) using Visiopharm® software (Medicon Valley, Denmark) using a 63x objective oil immersion lens. For each rat and RNAscope assay, 8 images spanning the entire rostro-caudal axis of the LC, ranging from −9.60 to −9.96 mm AP relative to Bregma [87], were analysed (**Figure 2A**).

**Figure 2:**
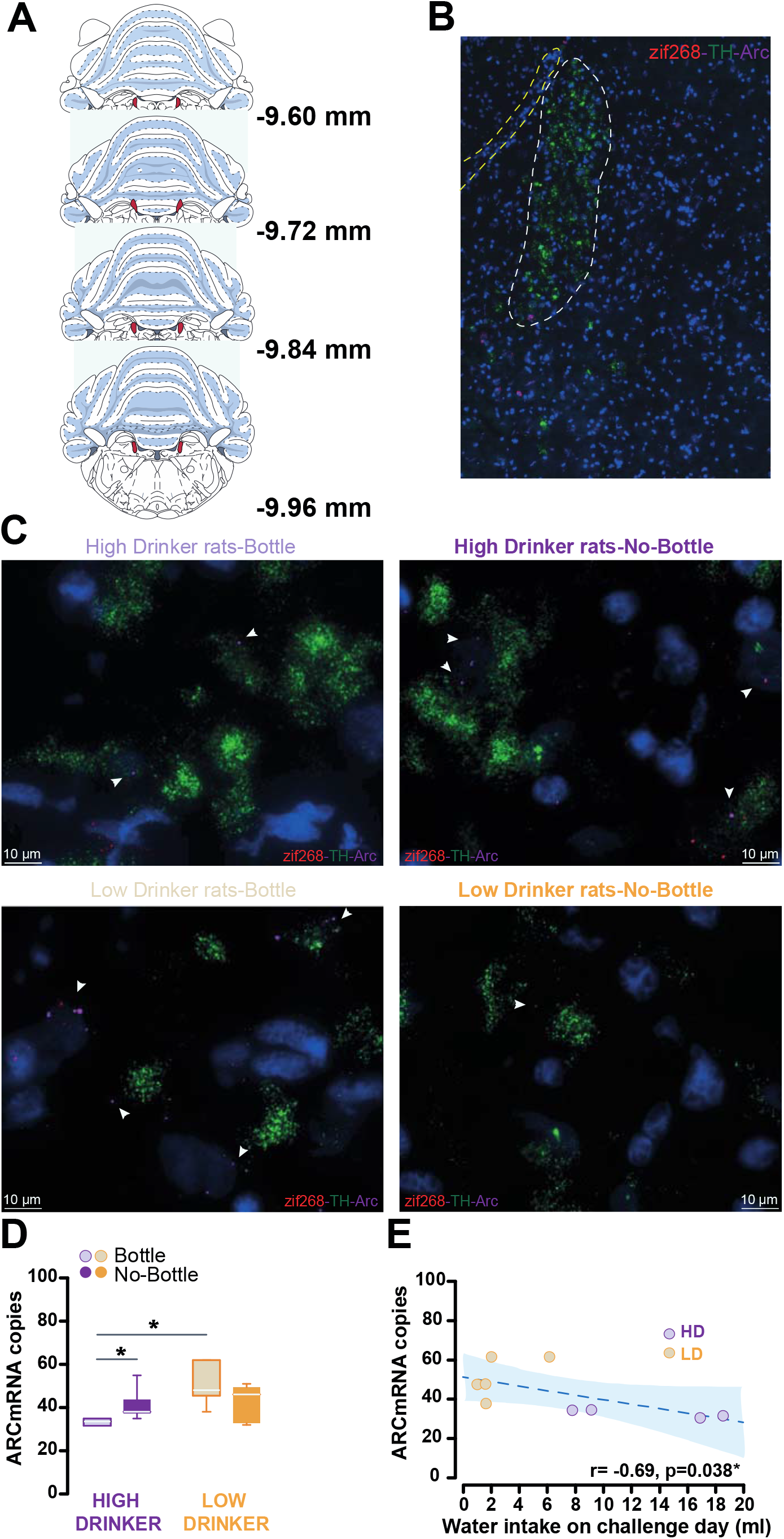
The expression of compulsive adjunctive behavior is associated with a decrease in Arc mRNA levels in the Locus Coeruleus. **A)** Schematic representation of the distribution of the coronal sections of the Locus Coeruleus (LC) ranging from −9.60 to −9.96 (relative to Bregma, coordinates shown next to each brain atlas template extracted from Paxinos & Watson [87]) that have been processed with RNAscope assays systematically to quantify the number of copies of target mRNAs. **B)** Representative superimage of a 12μm-thick coronal section, taken at 40X magnification showing the LC, where cell bodies of tyrosine hydroxylase expressing cells are located, delineated in white and a yellow outline of the edge of the fourth ventricle as anatomical reference, processed with RNAscope targeting the tyrosine hydroxylase, zif268 and Arc mRNAs. **C)** Representative 63x images of the LC from HD and LD rats that had, or not, the opportunity to express their adjunctive polydipsic behavior during the challenge session that preceded the harvesting of the brains, with the same green, red and magenta, fluorescent probes as in B) targeting TH, zif268 and Arc mRNA, respectively). The white arrows point the presence of Arc mRNA molecules. **D)** The individual tendency to eventually express compulsive adjunctive polydipsic drinking behavior was associated with a lower level of Arc mRNA expression in the LC in that HD rats showed fewer ARC mRNA copies in this monoaminergic nucleus than LD rats [Kruskal Wallis H(3)= 8.185, p= 0.043; HD-B vs LD-B p=0.027]. Preventing the animals from expressing their compulsive adjunctive polydipsic drinking behavior by the removal of the bottle resulted in an increase of Arc mRNA copies in the LC [Kruskal Wallis H(1)= 5.07, p= 0.0.243; HD-B vs HD-NB p=0.027], thereby suggesting that is the expression of compulsive polydipsic behavior which is responsible for the lower levels of Arc. **E)** At the population level, the number of Arc mRNA copies in the LC was found to be correlated with the level of polydipsic drinking during the challenge session [Spearman correlation, r= −0.692, p= 0.038].

Although RNAscope has recently gained popularity and is now widely used, the quantification strategy of the signal it yields has hitherto been sub-optimal, being limited to either the measurement of the signal intensity per channel/wavelength/probe or the quantification of mRNA positive cells, thereby missing out the opportunity to systematically quantify the number of mRNA molecules on a cell or a structure.

Several methods using semi-quantitative and quantitative analysis have been developed [88,89], but they too often rely on algorithms that require highly specialised coding skills, rendering them difficult to use by the many laboratories that do not have such expertise. In addition, most of the microscopy-based quantification methods and softwares available through different commercial platforms are semi-quantitative approaches that do not make full use of the single molecule quantification opportunity given by RNAscope. Thus, in order to maximize the information generated by the multiplex RNAscope assay, we developed a new analysis pipeline that goes beyond a simple quantification of light intensity to assess the levels of a target mRNA.

Our RNAscope signal analysis pipeline relies on a MATLAB (MATLAB - R2020a, The MathWorks Inc) script combined with a machine learning algorithm that uses ImageJ/FIJI (National Institutes of Health, Bethesda, MD, USA) to identify a single cell-delineated region of interest (ROI) based on a segmentation on the DAPI signal and the ensuing determination of an area (the ROI) around it within which to count single mRNA molecules.

mRNA molecules appear in images as bright “*dots*”, those with intensity exceeding chosen thresholds are considered true detections. For the detection of single mRNA molecules corresponding to the relevant channels 1, 2 and 4 the script enables the choice of independent threshold parameters (th1, th2 and th4). In order to remove background and maximise the signal-to-noise ratio, a difference of Gaussian filtering pre-processing step is applied. The raw image is blurred by convolution with a small Gaussian, σ1 = 0.5 pixels, to enhance signal. The background estimated by a larger Gaussian filtering, σ2 = 3 pixels, is subsequently subtracted from the image.

Nuclei segmentation is performed on the DAPI (third) channel. The pipeline allows the selection of a minimum area for nucleus detection. A Gaussian filter (σ=7 pixels) is applied to the DAPI image in order to smooth intensity inhomogeneities. Subsequently a multi threshold quantisation is performed, detecting three levels of nuclei intensities, since not all the DAPIs (or ROIs) are equally bright due to a different location along the Z-axis. Touching nuclei are separated via a watershed transformation (adapted to the levels of intensities detected in the image). Additionally, and as an optional step, all the DAPI segmentations can be manually corrected using the imageLabeler app in MATLAB.

However, watershed techniques are very susceptible to over-splitting and become less accurate in the case of elliptical cells and can be prone to segmentation errors if the nuclei are densely clustered together. In order to overcome these issues, we alternatively used StarDist [90] (in ImageJ/FIJI) for nuclei segmentation. StarDist is a segmentation method based on a deep learning, U-Net architecture that localizes cell nuclei approximated as star-convex polygons.

Once all the parameters have been set based on a sample image, and the detection of the different mRNAs and cell nuclei is deemed appropriate by the experimenter, they are kept unchanged for the entire RNAscope assay or batch image analysis

Finally, all the processed images pertaining to the same experiment are analysed, mRNA molecules closer than a certain distance (10 pixels) to a nucleus are assumed pertaining to that nucleus. The detections in each channel corresponding to each ROI are summarised and exported in Excel files.

The software package is available at https://gitlab.com/lemur01/rnascopeanalysis.

### Data and Statistical analyses

Data are presented as means ± SEM or box plots [medians ±25% (percentiles) and Min/Max as whiskers)] and analysed using STATISTICA 10 software (Statsoft, Palo Alto). Assumptions for normal distribution, homogeneity of variance and sphericity were confirmed using the Shapiro–Wilk, Levene, and Mauchly sphericity tests, respectively.

Water intake during the SIP procedure sessions was analysed using a repeated-measure analysis of variance (ANOVA) with sessions as within-subject factors and phenotype (High and Low drinkers; HD, LD) and Challenge (Bottle and no bottle; B, NB) as between-subject factor. Upon confirmation of significant main effects, differences were analysed using the Newman-Keuls post hoc test.

Ten representative animals of the HD and LD group were selected for RNAscope. The water intake of the last 3 SIP sessions was averaged and differences between the 2 groups (B, NB) independently for HD and LD were analysed using a Student’s t-test.

RNAscope images data belonging to the same animal were summed and then averaged across groups. The Kruskal Wallis test was used to quantify the different mRNA levels in the LC from HD and LD rats. Relationships between mRNA levels and water intake during SIP sessions were investigated using Spearman correlations and the p values obtained were subsequently corrected for multiple comparisons by the Benjamini-Hochberg method [91,92]. One animal belonging to the HD-B group was removed from the zif268-TH-Arc RNAscope data due to tissue damage during the assay. For all analyses, significance was set at α = 0.05. Effect sizes are reported as partial eta squared (pη^2^).

## Results

The cohort of forty-eight rats exposed to a SIP procedure developed a polydipsic adjunctive water drinking behavior over the course of 21 daily sessions [session *F*_20,320_ = 14.03, *p* ≤ 0.001, pη^2^= 0.46]. However, marked individual differences were observed in the tendency to develop compulsive adjunctive behavior, in that HD rats eventually drank more than 3 times as much water as LD rats by the end of training [main effect of group: *F*_1,16_ = 32.76, *p* ≤ 0.001, pη^2^= 0.67 and group x session interaction *F*_20,320_ = 8.01, *p ≤* 0.001, pη^2^= 0.33].

In line with previous reports [44,84], HD rats started to show excessive water intake on session 8, when their daily intake started to differ from their baseline and that of LD rats whose water consumption never differed from baseline (**Figure 1B**). The LD and HD individuals that were assigned to the bottle or no-bottle condition during the challenge session displayed similar levels of adjunctive drinking [t = 0.12 *p* = 0.90 and t = −0.22, *p* = 0.82, respectively] (**Figure 1C**).

Ten individuals, representative of the HD and LD groups, were selected for RNAscope assays which were deployed to measure Arc-TH-Egr1/zif268 and Arc-GFAP-Egr1/zif268 within the LC (**Figure 2A, 2B**).

The expression of compulsive adjunctive behavior was associated with a lower level of Arc mRNA expression in the LC. Thus, HD rats showed fewer Arc mRNA copies in the LC than LD rats [Kruskal Wallis H(3)= 8.185, p= 0.043; HD-B vs LD-B p=0.027] (**Figure 2C, 2D**) and the total number of Arc mRNA copies in the LC was negatively correlated to the intensity of polydipsic water drinking during the challenge session [Spearman correlation, r= −0.692, p= 0.038] (**Figure 2E**).

This lower level of Arc mRNA copies in the LC that characterised compulsive drinking in HD rats appeared to be specifically associated with the behavioral expression of the compulsion to engage in an adjunctive behavior as LC Arc mRNA levels were higher in HD rats that were prevented from expressing their polydipsic drinking by removal of the bottle during the challenge session [Kruskal Wallis H(1)= 5.07, p= 0.0.243; HD-B vs HD-NB p=0.027] (**Figure 2C, 2D**).

Further retrospective dimensional analyses revealed that the negative relationship observed between Arc mRNA copy number in the LC and the intensity of polydipsic water drinking observed during the challenge session in individuals that had access to the water bottle actually emerged on the 8^th^ SIP session, namely when HD rats started to develop excessive, compulsive adjunctive drinking leading them to differ from LD rats (**Figure 3**). This negative relationship was further shown to be primarily due to TH+ cells that did not co-express zif268 at the time of sacrifice (**Figure 3**). Thus, the percentage of Arc+ cells that were also TH+ but zif268-tracked the emergence of compulsive adjunctive drinking behavior following a pattern that was very similar, if not identical to that of the Arc mRNA copy number (**Figure 3**).

**Figure 3:**
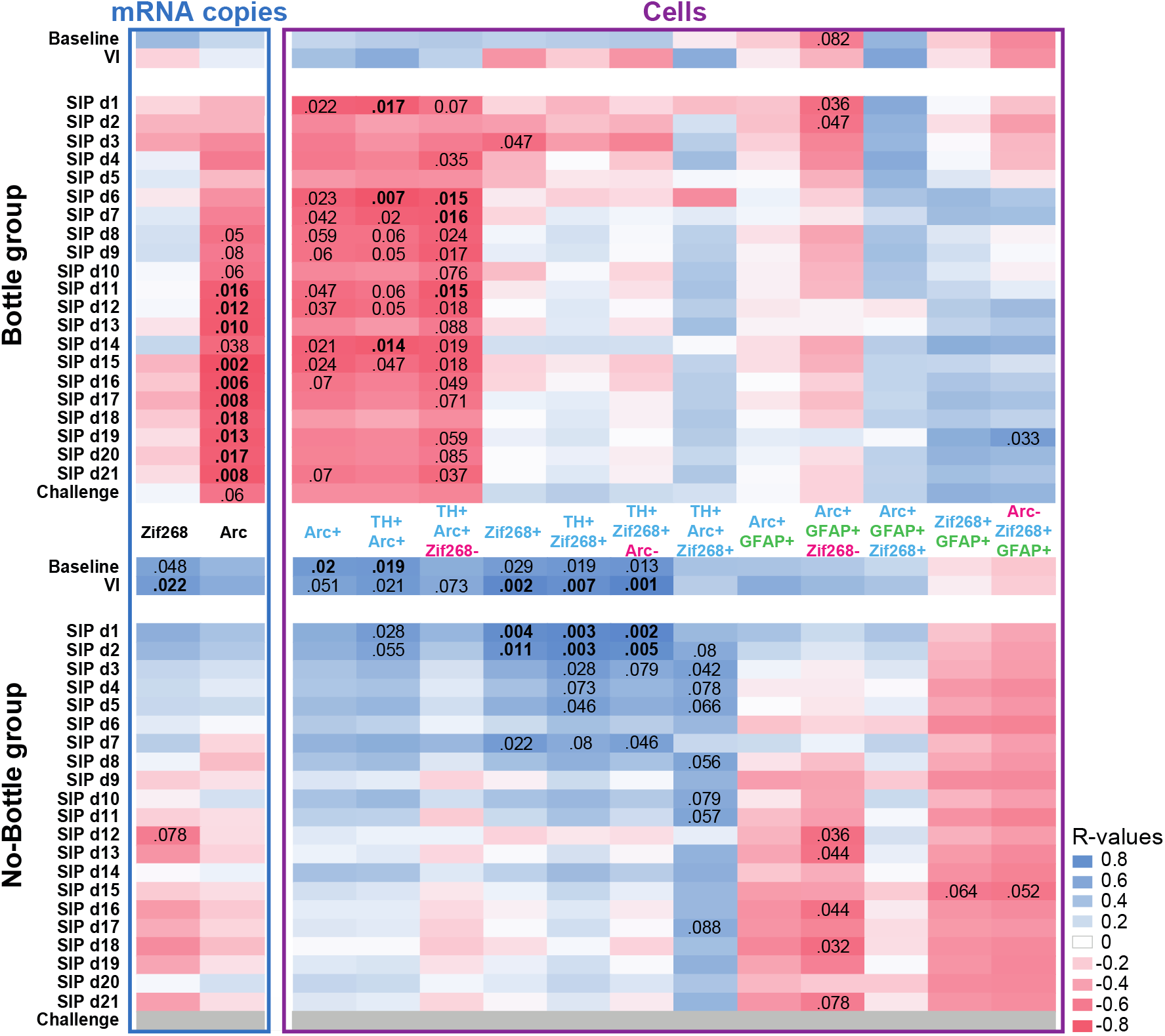
Downregulation of Arc mRNA levels in the LC, especially in TH+ but zif268-cells tracks the development of compulsive adjunctive polydipsic drinking behavior. The regulation of Arc mRNA levels within the LC by the expression of polydipsic drinking was not uniform across its different cell types and functional ensembles as shown by correlation matrices which track the relationship between the level of polydipsic drinking and (i) the total mRNA copies of zif268 and Arc in the LC and (ii) the representativity of various cell types characterised for their expression of the tyrosine hydroxylase, GFAP, zif268 and Arc, as well as any of their combinations. This dimensional analysis revealed that the negative relationship observed between the number of Arc mRNA copies in the LC and polydipsic drinking only emerged from session 8 onwards, exactly at the time when HD rats started to diverge from LD rats on their route towards the development of compulsive polydipsic drinking. This relationship was not observed for zif268, a distinction suggestive of a differential contribution of the two plasticity markers to the development of compulsive adjunctive behavior and supported by a very similar pattern of correlation between polydipsic drinking and the percentage number of cells expressing Arc, but not zif268. Further analysis of these matrices revealed that the relationship between polydipsic drinking and Arc mRNA levels was only present in neurons (no correlation was found in GFAP+ cells), more precisely in TH+ neurons and only in those that do not belong to a zif268+ ensemble. Removing the water bottle at test, which prevented rats from expressing their adjunctive polydipsic behavior, be it compulsive or not, disrupted the correlation with Arc mRNA levels, thereby revealing that the downregulation of Arc in TH+ zif268-cells ensembles track the expression of a compulsive adjunctive response.

The inverse relationship between Arc activation in the LC TH+ neurons and compulsivity was not observed for the immediate early gene zif268 or when rats did not have the opportunity to express their adjunctive behavior, be it compulsive or not, during the challenge session (**Figure 3**). What emerged in these individuals was a consistent positive relationship between the levels of drinking prior to the development of excessive adjunctive behavior in vulnerable rats and activation of neurons in the LC, as shown by consistent positive correlations between water intake on baseline and early SIP sessions and the percentage of zif268+ cells, irrespective of the co-marker they express.

Further analyses confirmed that these relationships between polydipsic water drinking behavior and Arc recruitment were specific to neurons as correlations were not observed consistently in GFAP+ cells (**Figure 3**)

## Discussion

The results of the present study show that the development of compulsive adjunctive polydipsic behavior under SIP is associated with a decrease in Arc mRNA levels especially, but not restricted to TH+zif268-neurons in the LC, a monoaminergic nucleus long suggested to be involved in stress [51] and, more recently, in coping strategies [50,53]. While the overall population of rats exposed to SIP developed a polydipsic response over time, in agreement with previous studies [46,93], individual differences emerged from the first week of training, with some individuals, namely HD rats, losing control over their coping response and developing very high levels of polydipsic drinking, the time course and magnitude of which is in line with that reported in previous studies from our laboratory and others [44,46,84,94]. These HD rats showed a decrease in their Arc mRNA levels in the LC when expressing their compulsive adjunctive response, since prevention of the expression of their compulsive drinking at test resulted in an increase in Arc mRNA levels which reached those of LD rats. This decrease in Arc mRNA levels at test tracked the emergence of compulsivity more that 16 days beforehand, as revealed by the emergence of a systematic negative correlation between Arc mRNA levels and daily water intake on session 8, the time at which HD rats started to differ from LD rats in their expression of polydipsic drinking. All together these data suggest that downregulation of the activity of the immediate early gene Arc in LC, and especially in TH+zif268-, neurons contributes to the loss of control over adjunctive polydipsic drinking, which underlies in HD rats the vulnerability to develop compulsive behaviors.

These observations are in line with evidence for functional alterations of the noradrenergic system in patients with compulsive disorders including genetic polymorphisms in the COMT gene [95–97], involved in the break-down of noradrenaline, elevated plasma levels of noradrenaline metabolites [97] and altered neuroendocrine responses to adrenergic drug challenges [98–100]. Similarly, chronic systemic administration of the noradrenaline reuptake blocker atomoxetine, a treatment that results in a decrease in spontaneous activity of LC neurons via α2-adrenoceptor stimulation in the LC [101] by chronically increased extracellular NA levels, prevents the development of SIP in highly impulsive rats [44]. Thereby this suggests that atomoxetine both stabilises an altered noradrenergic function in highly impulsive individuals [102] and prevents the recruitment of LC noradrenergic activity following exposure to SIP [103]. At the neural systems level, the influence of atomoxetine on impulse control has been shown to be mediated by the medial portion of the Nucleus Accumbens Shell (NAcS) in that bilateral intracerebral infusions of atomoxetine into the NAcS, but not into the Nucleus Accumbens Core (NAcC) replicated the effects its systemic administration on the level of premature responses in the 5-choice serial reaction time task [104]. Future research is warranted to determine whether the TH+zif268-LC neurons that display a specific decrease in Arc mRNA levels associated with the development of compulsive adjunctive behaviours are those that project to this area of the NAcS [104,105]. Indeed, when we performed unilateral infusions of a retrograde CAV2-GFP virus at different anterioposterior coordinates covering the entire rostro-caudal extent of the NAcS we observed that the injection site that resulted in the highest density of labelled putative neurons in the LC was that specific domain of the NAcS in which infusions of atomoxetine result in a decrease in impulsivity *(unpublished)*

Nevertheless, the results of the present study shed a new light on the cellular and molecular basis of the involvement of the LC in the development of compulsion resulting from the loss of control over coping strategies, one of the earliest evidence for which was the demonstration that bilateral lesions of the LC decrease water consumption in rats exposed to a SIP procedure without influencing homeostatic thirst [55]. Since, it has been shown that the individual tendency to engage in active vs passive coping responses and the ensuing differential resistance to stress engages different adaptations in the neural circuits controlling the LC-NA stress response system [50] in a genetically determined manner [54].

The mechanisms by which downregulation of Arc in TH+zif268-neuronal ensembles in the LC accompanies or perhaps underlies, the expression of compulsive adjunctive behaviors in vulnerable individuals remain unknown. Arc is broadly expressed at low levels under resting conditions and its transcription is rapidly and transiently induced following synaptic integration [106,107]. Arc expression is regulated by emotionally relevant experiences, including stressful situations [66,78] or alcohol withdrawal, in a large number of brain regions; including the BLA [108] and the NAc in which a large increase in Arc mRNA and protein levels is triggered by exposure to social defeat stress [66]. Viral overexpression of Arc in the NAc of Arc-knockdown mice is sufficient to rescue anxiety-like behaviors [109], thereby demonstrating causally that striatal Arc contributes to the regulation of anxiety [109], supposedly through its influence over dendritic plasticity [78]. Indeed, Arc mRNA is trafficked to neuronal dendrites [110] and, induced by neuronal activity [111], translated into proteins that can promote both synaptic strengthening and weakening [112]. In agreement with these observations, Arc has been shown to be involved in multiple forms of glutamatergic plasticity [113–115]. Overexpression of Arc blocks the homeostatic increase in AMPA-type glutamate receptors (AMPARs), whereas its decrement results in increased AMPAR function and a reduction of homeostatic scaling of AMPARs [116]. Thus, Arc is well positioned to influence glutamate-dependent plasticity and its function in learning, and memory [117]. In agreement with other clinical and preclinical studies having established the involvement of altered glutamatergic function in compulsive disorders, the tendency compulsively to express a polydipsic adjunctive behaviour under a SIP procedure has been shown to be associated with a lower level of glutamate in the medial prefrontal cortex [47], and reduced by glutamatergic drugs such as memantine [118]. Together with the previous evidence that compulsive symptoms are decreased by glutamatergic drugs in patients [119,120], this observation suggests that the downregulation of Arc mRNA levels in the LC observed in vulnerable individuals when they express a compulsive adjunctive behavior may be reflective of altered glutamatergic integration by LC neurons. Further research is warranted to establish which circuit, if any, among the glutamatergic inputs to the LC that include the paragigantocellularis nucleus, the lateral habenula and prefrontal cortex [121–123], are involved in these adaptations.

Within the LC, this study has revealed that a differential recruitment of Arc- vs zif268-functional ensembles is associated with resilience to the loss of control over coping strategies. Thus, rats which expressed a compulsive polydipsic response showed a specific downregulation of Arc mRNA in TH positive neurons that did not co-express zif268. In contrast, the mRNA levels of each marker in cellular ensembles or cell types in which they were both expressed did not correlate with polydipsic drinking when it became compulsive. While little is known about the functional relationship between Arc and zif268, the transcription of the former has been shown to be under direct regulation of Egr family of transcription factors to which the latter belongs [124], thereby suggesting that the two factors should show converging mRNA levels, as they do in ensembles in which their mRNA levels do not correlate with compulsive drinking. It has indeed been shown that exposure to nicotine, for instance, results in converging increases in Arc and zif268 mRNA levels [80]. However, such studies relied on classical in situ hybridisation, which did not enable a multiplex approach necessary to determine whether the increases in mRNA levels were occurring in the same cell type or across different ensembles. In other studies on cortical and hippocampal neurons, it has been revealed that while most of the cells that expressed Arc also expressed zif268, some zif268+ cells did not express Arc. This degree of independence between the functional recruitment of the two IEGs, which would be a prerequisite for the apparent higher sensitivity to behavioural demands that the effect IEG Arc shows as compared to other IEGs, including Zif268 [125] suggests that there may be regulatory mechanisms in addition to, and competing with those involving zif268 in the control of Arc mRNA levels. Transcriptional activation of Arc by zif268 is indeed completely inhibited by coregulatory factors such as Nab2 [126]. Further research will be necessary to better understand the molecular mechanisms that contribute to the emergence of this Arc+/TH+/zif268-specific LC neuronal ensemble in which a downregulation of Arc selectively characterises the expression of compulsive adjunctive behaviour in vulnerable individuals.

Altogether, the findings of the present study identify an ensemble in the LC characterised by a decrease in the expression of Arc in TH+zif268-neurons as a cellular marker of the expression of compulsive adjunctive drinking, thereby opening avenues for a mechanistic understanding of the role played by specific neuronal ensembles in the LC in coping behaviors and their compulsive manifestations.

## Acknowledgements

This work, carried at the department of Psychology of the University of Cambridge, was funded by a UKRI grant (MR/N02530X/1) to Barry Everitt, Trevor Robbins, Amy Milton, Jeff Dalley and David Belin and a Wellcome Trust Seed Award (109738/Z/15/Z) to DB. Leila Muresan was supported by a EPSRC grant (EP/R025398/1).

For the purpose of open access, the author has applied a Creative Commons Attribution (CC BY) licence to any Author Accepted Manuscript version arising.

## Authors contribution

CVS and DB designed the experiments. CVS carried-out the behavioral experiments and performed the histological procedures. LM wrote the MATLAB scripts for RNAscope quantification. CVS and DB performed the data analysis. CVS and DB wrote the manuscript.

